# SARS-CoV-2 amino acid substitutions widely spread in the human population are mainly located in highly conserved segments of the structural proteins

**DOI:** 10.1101/2020.05.16.099499

**Authors:** Martí Cortey, Yanli Li, Ivan Díaz, Hepzibar Clilverd, Laila Darwich, Enric Mateu

**Author notes:** Corresponding autor.

## Abstract

The *Severe acute respiratory syndrome coronavirus 2* (SARS-CoV-2) pandemic offers a unique opportunity to study the introduction and evolution of a pathogen into a completely naïve human population. We identified and analysed the amino acid mutations that gained prominence worldwide in the early months of the pandemic. Eight mutations have been identified along the viral genome, mostly located in conserved segments of the structural proteins and showing low variability among coronavirus, which indicated that they might have a functional impact. At the moment of writing this paper, these mutations present a varied success in the SARS-CoV-2 virus population; ranging from a change in the spike protein that becomes absolutely prevalent, two mutations in the nucleocapsid protein showing frequencies around 25%, to a mutation in the matrix protein that nearly fades out after reaching a frequency of 20%.

## 1. Introduction

The emergence of the novel *Severe acute respiratory syndrome coronavirus 2* (SARS-CoV-2) and the subsequent pandemic has become a health problem unparalleled in the last century. SARS-CoV-2 is thought to be originated from an animal coronavirus that successfully adapted to humans. The species of origin of SARS-CoV-2 has not been fully identified, but the virus seems to be related to SARS-CoV and other coronaviruses found in bats and other mammal species, although different from them (Chan et al. 2020; Lu et al. 2020; Zhou et al. 2020).

The SARS-CoV-2 genome size is around 30 kb with the typical gene structure known in other betacoronaviruses: starting from the 5′, more than two-thirds of the genome comprises orf1ab encoding polyproteins (nsp1 to nsp15), while the last third consists of genes encoding major structural proteins; including spike (S or ORF2), envelope (E or ORF4), membrane (M or ORF5), and nucleocapsid (N or ORF9) proteins. Additionally, the SARS-CoV-2 contains at least 6 minor structural proteins, encoded by ORF3a, ORF6, ORF7a, ORF7b, ORF8, and ORF10 genes (Khailany et al. 2020).

The first cases of the novel coronavirus associated disease (CoVID-19) have been traced to the Chinese province of Hubei in early December 2019 (https://www.who.int/csr/don/12-january-2020-novel-coronavirus-china/en/). Although the actual index case is not really known, the first sequence of the novel coronavirus was produced within weeks from the emergence of the disease (Zhu et al. 2019). As of the moment of writing this paper, more than 16,000 sequences have been produced in less than five months since the start of the pandemic. This is a unique opportunity to gain insight on the evolution of a betacoronavirus in a completely naïve human population. In this context, viral variants efficiently transmitted will have less influence of the selection exerted by the immune response, since most transmissions will occur from individuals before the development of an efficient immune response to naïve recipients.

The aim of the present study was to determine the amino acid substitutions in viral proteins that were widely present in available sequences of SARS-CoV-2, relating them to the known chronology of the pandemic. Also, the mutations found were assessed in order to try to understand its potential significance for viral fitness.

## 2. Material and methods

### 2.1. Sequences

SARS-CoV-2 sequences were retrieved from GISAID database (https://www.gisaid.org/). The full set of sequences used in the present study included the 12,562 high-quality complete sequences available on May 3^rd^, 2020. Additionally, a reference sequence from SARS-CoV, pangolin, civet, and three from bat coronaviruses (Genbank accession numbers AY278741, MT084071, AY572034, KY417146, MN996532 and MK211376, respectively) were used for comparative purposes. The set of sequences were arranged chronologically by date of isolation after the first reported SARS-CoV-2 sequences (identified as Wuhan-01 from December 24^th^, 2019).

### 2.2. Analysis of non-synonymous mutations and selection of mutations to be studied

Complete genomes were aligned using the multiple alignment program ClustalW (Thompson et al. 1994) and consequently split by week according to their isolation date with the sequence alignment editor Bioedit (Hall 1999). Using Wuhan-01 as the reference, an arbitrary date for the 1st report of the amino acid changes at the end of February 2020 was set to represent an early date of the pandemic, three months after the initial case was reported. Also, an arbitrary cut-off frequency of 10% was set to select the amino acid substitutions that were considered widespread. Thus, any substitution reported before the end of February and present in 10% or more of the frequencies in a given week was studied. In order to check if the variants identified presented a worldwide distribution, their geographical distribution was summarized by continents.

For each substitution, an alignment with the homologous proteins of SARS-CoV, civet, pangolin, and bat coronaviruses was performed to assess whether the mutation affected conserved or variable regions.

### 2.3 Comparison with predominant amino acid substitutions in early, mid and late cases of SARS epidemic of 2003-04

To compare predominant non-synonymous mutations occurring during different phases of the 2003 SARS-CoV epidemic, all sequences of SARS-CoV available at Genbank for which date of the case was available (directly or through literature search) were collected. Sequences were classified as early, mid or late based on the common classifications of cases (He et al. 2004). Analysis of mutations was done similarly to SARS-CoV-2.

### 2.4 Analysis of the potential biological significance of the observed substitutions

Modelling of the original S protein in Wuhan-01 and the mutant protein was produced using SWISS-MODEL protein template 6vsb.1.A (https://swissmodel.expasy.org/). Accuracy of the models was assessed by the Global Model Quality Estimation (GQME) and the Qualitative Model Energy Analysis (QMEAN) scores. 3D structures were rendered using PyMOL (The PyMOL Molecular Graphics System, Version 2.3.4. Schrödinger, LLC). The same program was used to determine changes in the protein structure or distances between atoms or residues. The set of proteomic utilities in EXPASY (https://www.expasy.org/proteomics) was used to check for different aspects on the mutant proteins (motifs, phosphorylation sites, etc.). PROVEAN 1.1. (http://provean.jcvi.org/, Choi et al. 2012) was used to gain insight on whether the mutation could be deleterious or neutral. Changes in the secondary structure of proteins were predicted by using the CFSSP: Chou & Fasman Secondary Structure Prediction Server (https://www.biogem.org/tool/chou-fasman/, Ashok 2013).

When mutations affected known epitopes, impact on antigenicity was evaluated by means of epitope prediction tools in the IBDB Resource web (http://tools.iedb.org/).

## 3 Results

### 3.1. Amino acid substitutions with significant spread in the population were mainly located in conserved segments of structural proteins of SARS-CoV-2

The analysis of the set of sequences revealed that only 8 amino acid substitutions across the viral genome appeared before the end of February 2020 and gained prevalence over 10% of the known isolates at a given time point, measuring time in weeks after the first available sequence (Fig. 1). Of these, 7 substitutions were in the structural proteins (ORF2-S, ORF3a, ORF5-M, ORF8, and ORF9-N) and one in the ORF1ab, specifically in nsp6. Concerning their location and date, four appeared in China during January. The other four appeared in Europe during the second fortnight of February, but within a week, they were also reported in other continents (Supplementary Table S1). Interestingly, the only substitution that became fully predominant was Asp614Gly in the spike protein (ORF2-S). Gly57His in ORF3a reached a frequency of 50% at the moment of writing this paper. It is worth noting that when the sequences where analysed by continents, all mutations were spread worldwide, except the 175Met in the ORF5-M, that was absent in Africa (Supplementary Table S1).

**Figure 1.**
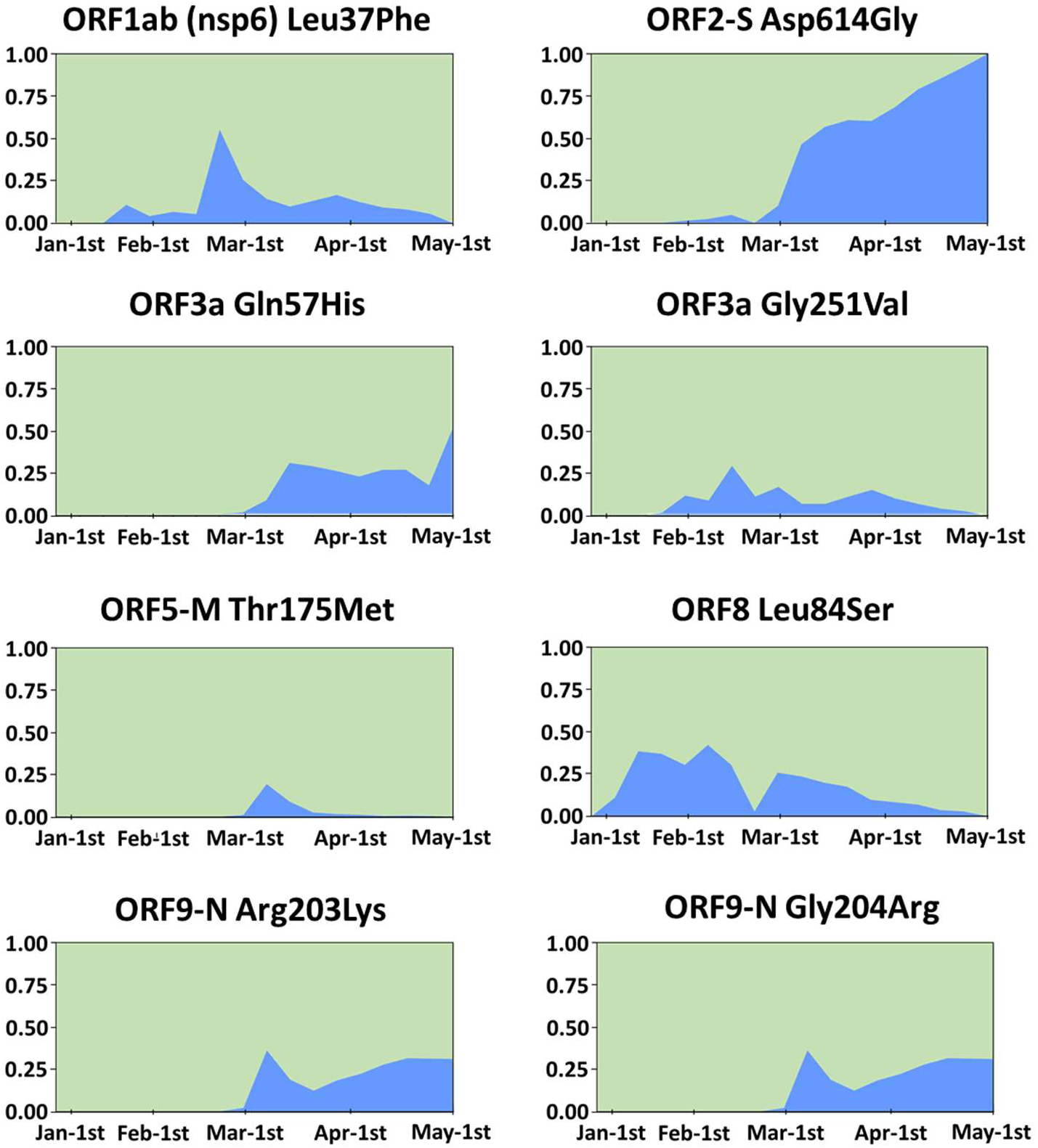
Temporal trends in the emergence and prevalence of the amino acid substitutions that appeared before the end of February 2020 and were present in >10% of SARS-CoV-2 sequences at any time between December 2019 and May 2020. The X-axis represents time measured as weeks since December 24^th^, 2019 and the Y-axis represents the proportion of known sequences harbouring a given mutation.

Next, we examined whether these substitutions were located in variable or conserved regions of the viral genome. Interestingly, most of them corresponded to residues that were conserved in SARS and related betacoronaviruses of pangolins, civets, or bats (Fig. 2).

**Figure 2.**
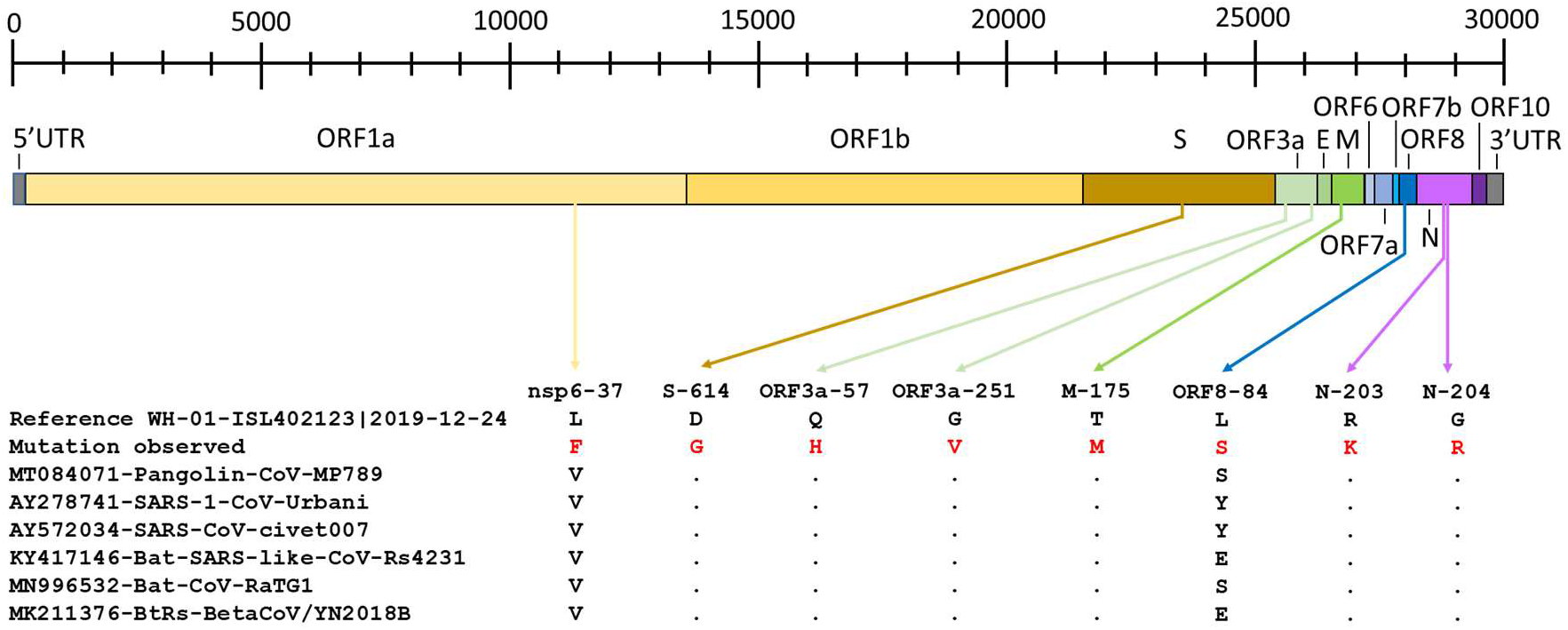
Location of the mutations found in the present study and corresponding amino acids in pangolin, SARS-CoV, civet, and bat-related coronaviruses. One-letter code is used to represent amino acids. A dot is used to indicate a conserved residue compared to the first sequence in the alignment (SARS-CoV-2 isolate Wuhan-01).

The comparison with non-synonymous mutations that gained predominance in SARS-CoV showed a different pattern. From early to late phases of the SARS epidemic of 2003, 11 substitutions gained wide spread. Three of them were located in nsp3, four in nsp4, one in nsp16, and three in the spike protein (Supplementary Table S2). However, mutations in SARS-CoV were located in conserved positions of civet and bat-related coronaviruses but, those positions were different in pangolin.

### 3.2. Substitutions in the major structural proteins S, N and M

The examination of the spike protein sequences of SARS-CoV-2 revealed that the Asp614Gly mutation that appeared in January 2020 in Shanghai, gained predominance with time. Thus, by the end of April 2020 it was almost 100% prevalent (Fig. 1).

Next, we checked whether this mutation emerged in a particular SARS-CoV-2 clade or its spread was irrespective of the clade where a given isolate could be allocated. The Gly614 could be found in different branches across the phylogenetic tree of SARS-CoV-2 (available at https://nextstrain.org/).

Residue 614 is located in the S1 domain of the spike protein. Modelling of the original and mutant proteins using template 6vsb.1.A from SwissModel of the S protein of SARS-CoV-2 (Wrap et al. 2020) showed that by changing Asp614 by Gly614, the distance to Thr859 and its side chain increased (from 2.7 to 6.2 Å), creating a cavity and a more relaxed structure (Fig. 3). The analysis of the antigenicity of the potential epitope homologous to 597-625 of SARS-CoV did not show any significant difference between mutants.

**Figure 3.**
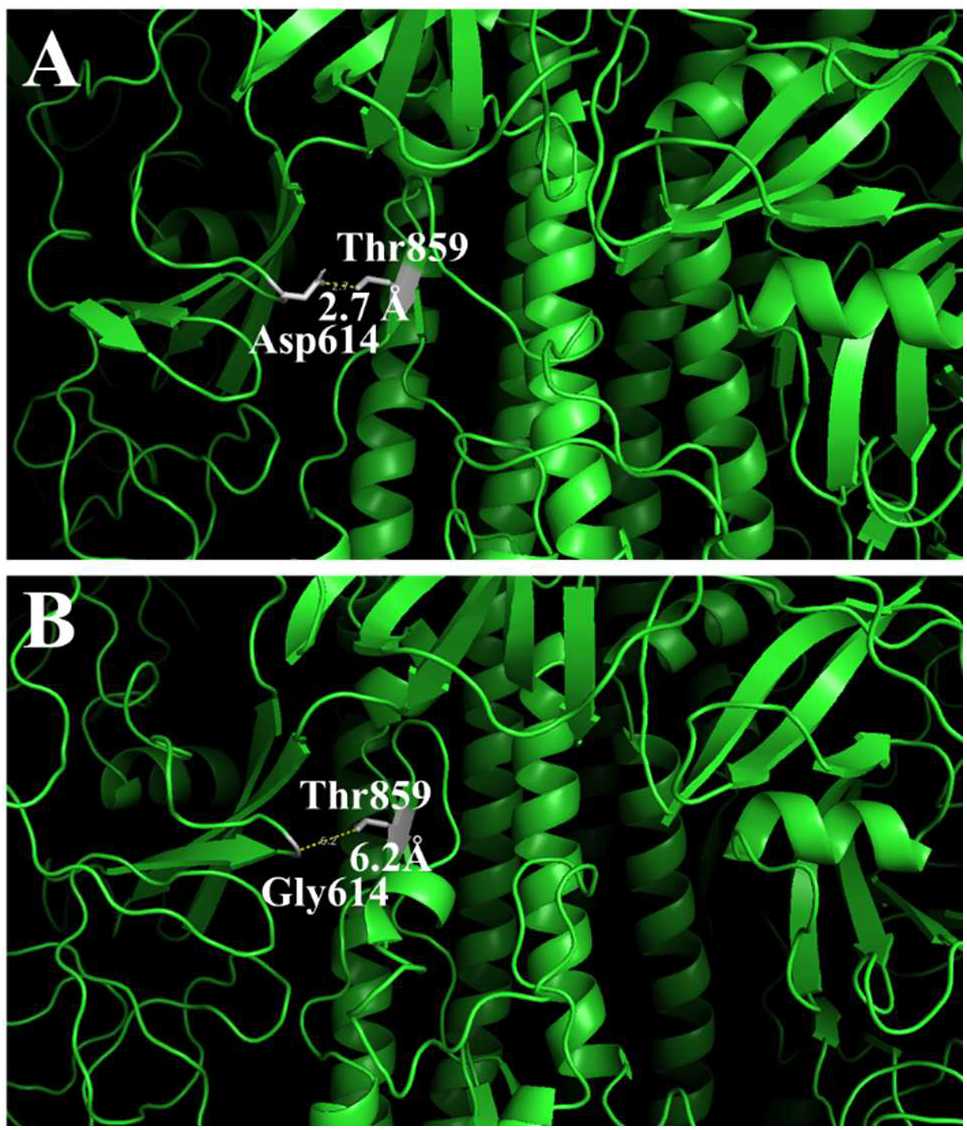
Changes in the 3-D structure of the S-protein in the original (A) and mutant (B) proteins. The pictures show amino acid residues on 20Å distance from Asp614 (A) or Gly614 (B) and their distance to Thr859.

In the nucleocapsid phosphoprotein, a double mutation (Arg203Lys-Gly204Arg) was observed to gain predominance during the pandemic (Fig. 1). This mutation is in the serin-rich (SR) segment of the linker region (LKR) of the protein. The N protein of SARS-CoV is known to be bound by Ubc9, a ubiquitin conjugating enzyme of the sumoylation system, probably at the SR segment. Since lysines are targets for sumoylation, we next checked whether this substitution could introduce a sumoylation motif. The prediction using two different methods JASSA v4 (http://www.jassa.fr/index.php?m=jassa) and GPS-SUMO (http://sumosp.biocuckoo.org/) was negative. Predicted phosphorylation sites and enzymes (NetPhos 3.1.) were not significantly different between variants. The observed mutations were predicted to be neutral by Provean.

In the matrix protein, the main change was the substitution of Thr175 by Met175. Thr175 was predicted to form part of a motif known to interact with 14-3-3 proteins and of a potential phosphorylation site (173-SRTLSYYKL-181) targeted by protein kinases A (PKA) and C (PKC), ribosomal s6 kinase (RKS), and DNA-dependent protein kinases. The substitution of the THR by a MET was predicted again to be a phosphorylation site (for PKA, PKC and RKS). Provean predicted that the introduction of M175 was deleterious (score −3.135). Since no reliable model was available, 3D structure could not be assessed. Interestingly, the presence of a M175 in the M protein occurred together with the 203KR204 mutation in 98% of the cases (p<0.0001). The rapid decay of this mutation (Fig. 1) would be consistent with a deleterious effect.

### 3.3. Substitutions in the minor structural proteins 3a and 8, and the non-structural protein nsp6

Two substitutions were identified in the protein encoded by ORF3a. The first was a substitution of Gln57 present in the Wuhan-01 isolate by His57, and the second was the substitution of Gly251 by Val251. To note, only 0.04% of the sequences harboured the double His57/Val251 mutation. The presence of His in residue 57 would be expected to result in an increased positive charge at that site.

The Gly251Val mutation occurred in a predicted serine-phosphorylation site 248-TID(**G/V**)<u>S</u>SGVV-256. The introduction of a Val reduced the prediction scores for the site from a maximum >0.90 with Gly for phosphokinase B and ATM serine/threonine phosphokinase to 0.73-0.82 for the same enzymes. It is worth to note that the amino acid at this position was strongly correlated with the amino acid present in position 57 of the same protein. Thus, among the sequences harbouring His57 in ORF3a, 99.8% were associated with Gly251 and only 0.2% with Val251. In contrast, for Gln57, 17% of the sequences harboured Val251 (p<0.001). Similarly, His57 was only found in sequences harbouring Arg203-Gly204 in the N protein, while Gln57 was found simultaneously with Arg203-Gly204 (70% of the cases) or Lys203-Arg204 (30%) (p<0.001). Both mutations were predicted as deleterious (scores of −3.286 for His57 and −8.581 for Val251).

The lack of a reliable model made impossible to make any prediction of the impact of those mutations on the 3D structure or the interactions between residues. However, the checking of the potential in the secondary structure of the protein revealed that mutation Gln57His resulted in the elimination of a turn of the protein predicted to be in Ser58. Similarly, mutation Gly251Val eliminated the turn at Gly251 but did not affect the turn predicted for Ser253 (Supplementary Fig. S3).

Regarding the ORF8 protein, it is worth noting that the mutation of residue 84 (Leu84Ser) happened simultaneously with a silent mutation in nucleotide position 8987 (ORF1ab, nsp4, A→T). This permitted to distinguish 2 clades in the initial weeks of the pandemic that contained isolates from Wuhan, Shanghai, and Hong-Kong (Supplementary Fig. S4). These clades did correspond to the L and S types reported by Tang et al. (2020). This mutation was predicted to be neutral.

Finally, the last change was found in the nsp6 protein, Leu37Phe, which significance was unclear. This mutation was also predicted to be neutral.

## 4. Discussion

The present SARS-CoV-2 pandemic is a worst-case scenario of the introduction of a new agent that transmits easily in a completely naïve population. In this context, transmission events occur in an uncommonly high scale, in a very short period of time and with little selective pressures from the immune system if compared to an endemic situation. This scenario would permit the arising of a great diversity of viral variants of which the fitter could be expected to gain predominance.

In the present study, we identified 8 early mutations in the SARS-CoV-2 genome that gained prevalence over 10% at some point during the pandemic. This cut-off was arbitrarily set to discriminate random mutations and errors in sequencing from changes that might have a bigger impact. Certainly, this approach has the limitation of neglecting some mutations with lesser prevalence that still can be biologically significant. Time will show it.

It is worth noting that 7 out of 8 of the widely spread mutations occurred in residues that were highly conserved in related coronaviruses of bats, pangolins, civets, or in SARS-CoV (Fig. 2). Conserved regions are usually assumed to be functionally relevant and thus, mutations in them may have deleterious effects or can be hardly tolerated; if so, they will be probably removed in the future. A mutation in a highly conserved region that becomes widespread and persists can be thought as representative of a change that increases viral fitness. In the present case, we found three different situations: mutations that expanded and rise to predominance, mutations that expanded to a certain extent and fade out, and mutations that are apparently expanding but not yet predominant. This pattern affecting conserved regions was also seen for SARS-CoV although the affected proteins were different. Interestingly, in SARS-CoV-2 most of the mutations were in structural proteins, while in SARS-CoV were in non-structural ones, suggesting that the adaption process from the original host species to human was different in these two cases. When the spike protein was examined, this difference was more obvious. Mutations in SARS-CoV occurred in positions conserved in the civet and bat-related coronaviruses but different from those of pangolin and SARS-CoV-2. In contrast, spike mutation in position 614 of SARS-CoV-2 affected a residue that was conserved in betacoronaviruses of pangolins, civets, bats, and SARS-CoV. This would be compatible with a scenario where those mutations affected the viral fitness for that particular new host, namely humans. The scale of the viral replication in the scenario of a pandemic would be unprecedented for those coronaviruses and will provide the probability for such beneficial mutations to appear and expand.

The Asp614Gly mutation in S protein is an example of a mutation becoming fully predominant. A previous report (Korber et al. 2020) already indicated its emergence. Recently, Bhattacharyya et al. (2020) suggested that the predominance of this mutation as the pandemic advanced and the low proportion in initial phases of it was related to a single nucleotide deletion in the transmembrane protease serine 2 (TMPRSS2) that is common in Europeans and North Americans but rare in East Asians. The Asp614Gly mutation would introduce a cleavage site for that enzyme. This would explain, at least partially, the spread of this mutant outside Asia.

We do agree with Korber et al. (2020) regarding the possibility that the Asp214Gly mutation produced a laxer interaction between S1 and S2 spike domains that might facilitate shedding of S1 from membrane-bound S2. In SARS-CoV, the segment of S protein including residues 597-625 contains epitopes inducing both neutralizing antibodies (positions 604-625 in SARS-CoV) and antibodies participating in an antibody-dependent enhancement (ADE) in animal models (residues 597-625) (Wang et al. 2016). The mutation Asp614Gly would affect the epitope segment inducing ADE but not the one inducing neutralizing antibodies. Although, it has been hypothesized that ADE may have a role in CoVID-19 (Tetro 2020) this has not been demonstrated (Sharma 2020). The effects of the mutation on the immune escape or the transmission potential cannot be concluded at this moment.

Regarding the mutations found in the nucleocapsid phosphoprotein, the first surprising fact was to find two consecutive substitutions in the highly conserved serine-rich segment of the LKR region of the protein. The LKR region is essential for conferring flexibility to the protein as well as for cell signalling, binding to RNA and to M protein. It also contains multiple phosphorylation sites in the SR segment that are thought to be essential (Reviewed by McBride et al. 2014). Since the mutant nucleocapsid introduced a Lys residue - a canonical target for sumoylation - and sumoylation occurs in this region by analogy to SARS-CoV (Fan et al. 2006) we also tested it. No significant differences were determined between the original and the mutant sequences using prediction tools for phosphorylation or sumoylation.

Interaction of the nucleocapsid protein with the M protein is thought to happen between the SR-region and the C-terminal domain of M (Escors et al. 2001; Kuo and Masters 2002). We found that the M175 phenotype of the M protein was almost exclusively associated to the 203K-204R phenotype of the nucleocapsid protein. It is tempting to hypothesize that the above-mentioned residues may be involved in such interaction. Besides this, the introduction of an additional charge in the SR segment may enhance interaction with RNA which core is negatively charged.

Changes in the ORF3a protein have been recently reported to define microclonal clades of SARS-CoV2 (Issa et al. 2020). We have found that the introduction of Val251 was apparently non compatible with His57. We determined that, probably, such mutations affected the secondary structure by changing the number of turns. Considering that both residues are out of the functional domains proposed by Issa et al. (2020), those changes in the structure of the protein may modify interactions enough to be non-compatible. Interestingly, mutations in ORF3a protein and mutations in the nucleocapsid protein were related. To our knowledge, these two proteins have not been investigated for interactions; however, this finding suggests that might be an interaction between them.

ORF8 mutation Leu84Ser was reported before (Tang et al. 2020). The authors suggested that the Leu variant (called L) is more aggressive and spreads easier. The evolution of the proportion of strains harbouring this mutation would not support the idea of a higher transmissibility of the L variant since its frequency clearly declined. Certainly, the introduction of such variant in different countries applying control measures earlier or later could have had an impact on the spread of the variants as well.

Finally, the nsp6 mutation at position 37 is difficult to interpret. The nsp6 of coronaviruses is part of the replication machinery of the virus and has been reported to induce autophagosomes (Benvenuto et al. 2020). The same authors suggested that it may lead to a lower stability of the protein structure. Lacking a verified model for the protein it is difficult to assess whether this happens or not. Interestingly, the same authors indicated that the distribution of the Leu phenotype was restricted to Asia while the Phe was common in other parts of the world. According to our analysis, the Phe phenotype has almost disappeared in current sequences. The discrepancy could be originated in the fact that Benvenuto et al. (2020) analysed the 351 sequences available in a past moment while in the present analysis thousands of sequences from all over the world have been included.

In summary, the present is a comprehensive report of the amino acid mutations that gained spread during the SARS-CoV-2 pandemic up to now. Most of the substitutions gaining wide diffusion occurred in conserved positions indicating that they probably had a functional impact but, differently from SARS-CoV, they accumulated in structural proteins. Interestingly, most of these mutations faded out, except for the Asp614Gly in the S protein that became predominant suggesting that it contributed to viral fitness. Some others are still increasing in prevalence, like the mutations in the nucleocapsid protein here reported and that might be related to the mutations in the M protein. This data may serve to gain further insight in the evolution of SARS-CoV-2.

## 5. Acknowledgements

The authors would like to thank to all those who contribute in the fight against the SARS-CoV-2. We would like to thank as well all those who kindly shared genome data in publicly available databases, available for download from https://www.gisaid.org/.

## 6. Author contributions

MC, YL, ID, HC, LD and EM participated in all tasks of the present work.

## 7. Funding

Martí Cortey was funded by the Ramon y Cajal program (reference RYC-2015-17154). Hepzibar Clilverd was supported by a fellowship of the Spanish Ministry of Science and Innovation (program FPU). No other funding sources.

## 8. Conflict of interest

None declared.

## 9. Supplementary Material

**Supplementary Table S1.** Distribution by continents, in percentages, of the 8 amino acid mutations detected across the 12,562 SARS-CoV-2 genomes analysed.

| Location | nsp6-37 |  | S-614 |  | ORF3a-57 |  | ORF3a-251 |  | M-175 |  | ORF8-84 |  | N-203 |  | N204 |  |
| --- | --- | --- | --- | --- | --- | --- | --- | --- | --- | --- | --- | --- | --- | --- | --- | --- |
| **1st Report** | Wuhan 18/01 |  | Shanghai 23/01 |  | France 21/02 |  | Hong-Kong 23/01 |  | Netherlands 24/02 |  | Wuhan 05/01 |  | England 23/02 |  | England 23/02 |  |
| **Aa Change** | Leu | Phe | Asp | Gly | Gln | His | Gly | Val | Thr | Met | Leu | Ser | Arg | Lys | Gly | Arg |
| **Asia (n=741)** | 81,8 | 18,2 | 69,1 | 30,9 | 82,8 | 17,2 | 90,6 | 9,4 | 98,6 | 1,4 | 80,6 | 19,4 | 94,4 | 5,6 | 94,4 | 5,6 |
| **Europe (n=7,571)** | 81,1 | 18,9 | 20,6 | 79,4 | 83,3 | 16,7 | 86,1 | 13,9 | 92,0 | 8,0 | 93,2 | 6,8 | 68,6 | 31,4 | 68,6 | 31,4 |
| **America (n=3,238)** | 95,8 | 4,2 | 37,2 | 62,8 | 48,6 | 51,4 | 97,9 | 2,1 | 99,4 | 0,6 | 72,0 | 28,0 | 96,1 | 3,9 | 96,1 | 3,9 |
| **Africa (n=110)** | 90,3 | 9,7 | 14,7 | 85,3 | 80,0 | 20,0 | 96,8 | 3,2 | 100,0 | 0,0 | 92,6 | 7,4 | 90,4 | 9,6 | 90,4 | 9,6 |
| **Oceania (n=902)** | 69,7 | 30,3 | 46,5 | 53,5 | 69,9 | 30,1 | 87,4 | 12,6 | 95,5 | 4,5 | 79,5 | 20,5 | 86,7 | 13,3 | 86,7 | 13,3 |
| **Mean (n=12,562)** | **84,2** | **15,8** | **29,7** | **70,3** | **73,1** | **26,9** | **89,7** | **10,3** | **94,6** | **5,4** | **86,1** | **13,9** | **78,7** | **21,3** | **78,7** | **21,3** |

**Supplementary Table S2.** Localizations of the amino acid mutations in SARS CoV that gained prevalence compared with SARS-CoV-2, civet, pangolin, and bat-related coronaviruses.

| Protein | ORF1ab<br>(nsp3) | ORF1ab<br>(nsp3) | ORF1ab<br>(nsp3) | ORF1ab<br>(nsp4) | ORF1ab<br>(nsp4) | ORF1ab<br>(nsp4) | ORF1ab<br>(nsp4) | ORF1ab<br>(nsp16) | ORF2S | ORF2S | ORF2S |
| --- | --- | --- | --- | --- | --- | --- | --- | --- | --- | --- | --- |
| Aa position in the protein | 303 | 844 | 1298 | 6 | 231 | 307 | 332 | 138 | 242 | 347 | 1166 |
| Original SARS-CoV | T | I | F | W | A | A | A | K | L | R | E |
| Mutated Aa in SARS-CoV | I | L | L | C | V | V | V | R | S | K | K |
| AY572034-SARS-CoV-civet007 | T | I | L | W | A | A | A | R | S | R | E |
| SARS-CoV-2 WH-01-ISL402123 | T | I | A | W | A | A | V | K | T | R | K |
| MT084071-Pangolin-CoV-MP789 | T | I | Del | W | A | A | V | K | T | R | K |
| KY417146-Bat-SARS-like-CoV-Rs4231 | T | I | L | W | A | A | A | K | L | R | K |
| MN996532-Bat-CoV-RaTG13 | T | I | A | W | A | A | A | K | T | R | K |
| MK211376-BtRs-BetaCoV/YN2018B | T | I | L | W | A | A | A | K | P | R | K |

**Supplementary Figure S3.**
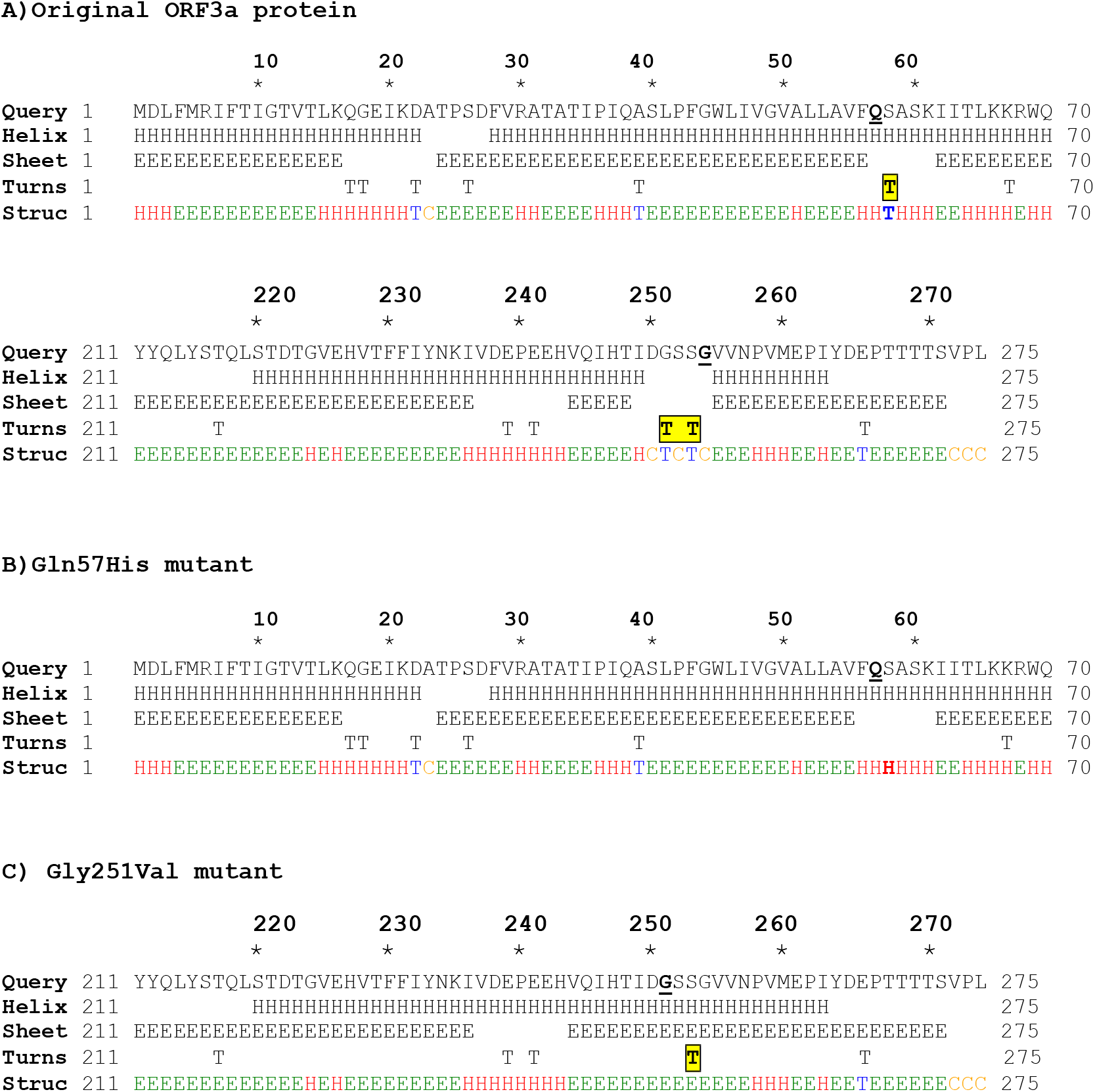
Secondary structure of the original ORF3a (A) or the mutant variants (B & C). H = alpha-helix, E = Beta sheet, T = Turn

**Supplementary Figure S4.**
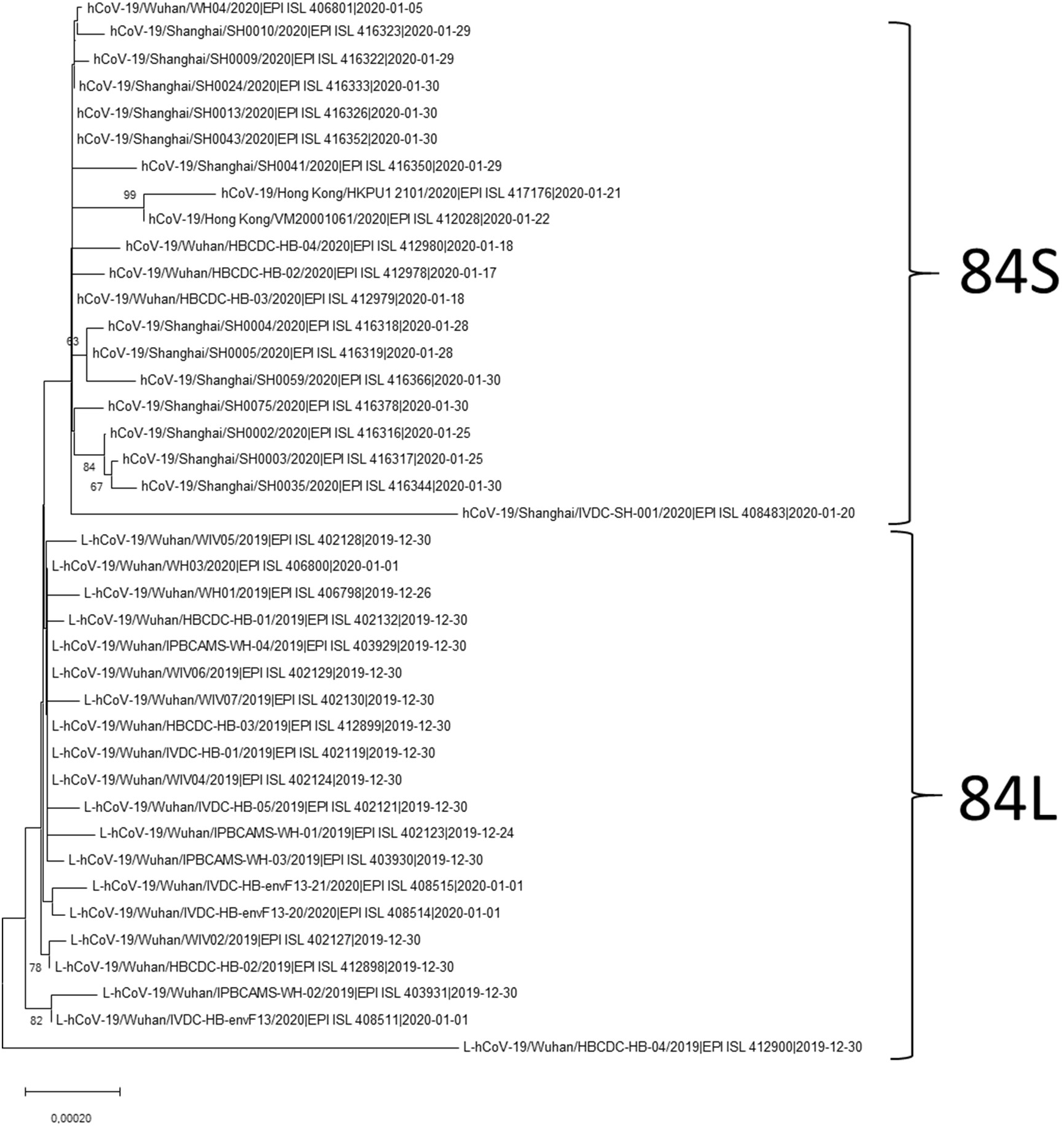
Neighbor-Joining tree based on the Tamura-Nei distances, constructed with the 20 earliest complete genomes harbouring a Leucine or a Serine in the amino acid residue 84 of the ORF8 protein. Isolation dates as reported in the GISAID database (https://www.gisaid.org/). Numbers along the internal branches represent their confidence after the initial dataset was resampled with 1,000 bootstrap replicates.

## Notes

### Competing Interest Statement

The authors have declared no competing interest.

